# Recovery of hand function after stroke: separable systems for finger strength and control

**DOI:** 10.1101/079269

**Authors:** Jing Xu, Naveed Ejaz, Benjamin Hertler, Meret Branscheidt, Mario Widmer, Andreia V. Faria, Michelle Harran, Juan C. Cortes, Nathan Kim, Pablo A. Celnik, Tomoko Kitago, Andreas R. Luft, John W. Krakauer, Jörn Diedrichsen

**Author notes:** These authors contributed equally to this work (Jing Xu, Naveed Ejaz). **Contact information** Jing Xu Department of Neurology, School of Medicine, Johns Hopkins University, Pathology 2-210 600 N. Wolfe St. Baltimore, MD 21287 USA.

## Abstract

Loss of hand function after stroke is a major cause of long-term disability. Hand function can be partitioned into strength and independent control of fingers (individuation). Here we developed a novel paradigm, which independently quantifies these two aspects of hand function, to track hand recovery in 54 patients with hemiparesis over the first year after their stroke. Most recovery of both strength and individuation occurred in the first three months after stroke. Improvement in strength and individuation were tightly correlated up to a strength level of approximately 60% of the unaffected side. Beyond this threshold, further gains in strength were not accompanied by improvements in individuation. Any observed improvements in individuation beyond the 60% threshold were attributable instead to a second independent stable factor. Lesion analysis revealed that damage to the hand area in motor cortex and the corticospinal tract (CST) correlated more with individuation than with strength. CST involvement correlated with individuation even after factoring out the strength-individuation correlation. The most parsimonious explanation for these behavioral and lesion-based findings is that most strength recovery, along with some individuation, can be attributed to descending systems other than the CST, whereas further recovery of individuation is CST dependent.

## Introduction

Human hand function comprises at least two complementary aspects: strength as manifest in a power grip, and control of individual finger movements as in piano playing. The most common observation after stroke is that both are impaired (Kamper and Rymer, 2001; Lang and Schieber, 2003). Weakness presents as difficulties in voluntarily opening of the hand, extending the wrist and fingers against resistance, and producing a strong grip (Colebatch and Gandevia, 1989; Kamper *et al.*, 2003). Loss of finger control manifests as inability to either move a single finger while keeping the others immobile, or to make complex hand gestures, both of which impair the ability to perform tasks such as typing or buttoning a shirt (Kamper and Rymer, 2001; Li *et al.*, 2003; Lang and Schieber, 2004). When strength does recover after stroke, control often remains impaired, causing lasting disability (Heller *et al.*, 1987; Sunderland *et al.*, 1989). However, the relationship between strength and control after stroke remains poorly understood. Separating the effect of stroke on finger strength versus control is a challenge given that most current clinical measurements conflate weakness with deficits in control. In the current study we therefore sought to develop a new paradigm that could measure these two aspects of hand function separately, and to investigate the relationship between strength and control over the time course of hand recovery after stroke. We were specifically interested to test whether these two components recover in a lawful relationship with each other, or whether they recover independently.

Existing behavioral tasks used to assess hand function after stroke, such as the Fugl-Meyer Assessment (FMA) (Fugl-Meyer *et al.*, 1975), the Nine-Hole Peg Task (9NPT) (Sharpless, 1982), and the Action Reach Arm Test (ARAT) (Lyden and Lau, 1991), are not designed to separate deficits in strength and control. To isolate these two aspects of hand function it is necessary to remove any obligatory relationship between them (Reinkensmeyer *et al.*, 1992), i.e. derive a control measure that is independent of strength. Intuitively, a rock climber may have stronger fingers than a pianist, but not necessarily superior control of individual fingers.

Schieber (1991) devised an individuation task that requires participants to move individual fingers while keeping the non-moving ones stationary. Movements of the passive fingers were used as a measure of loss of control. This paradigm however does not directly track the force relationship between active and passive fingers. In the paradigm used here, we first measured the maximum voluntary contraction force (MVF) that a participant could produce with each finger. We then asked participants to produce isometric forces over four sub-maximal levels with each finger, while keeping the passive fingers immobile. Even controls show involuntary force production (enslaving) on the passive fingers, which increases with the required active force level (Li *et al.*, 1998; Zatsiorsky *et al.*, 2000). The slope of the function of passive finger enslaving on active force thus provides a measure of individuation that is independent of strength.

Using this paradigm we tracked the recovery of hand strength and finger individuation in patients over a one-year period after stroke. One possibility is that strength and control recover independently. For example, a patient may remain quite weak but have good recovery of individuation, or a patient may recover a significant amount of grip strength but fail to individuate the digits. Alternatively, recovery may be such that when strength recovers so does individuation, because either they share a common neural substrate or repair processes are proceeding in parallel in separate neural substrates. Finally, lesion analysis allowed us to investigate whether there is any identifiable anatomical basis for any observed dissociation between strength and control deficits.

## Materials and Methods

### Participants

Fifty-four patients with first-time ischemic stroke and hemiparesis (34 male, 20 female; mean age 57.4±14.9 years) were recruited from three centers: The Johns Hopkins Hospital and Affiliates, Columbia University Medical Center, and The University Hospital of Zurich and Cereneo Center for Neurology and Rehabilitation. According to the Edinburgh Handedness Inventory (Oldfield, 1971), Forty-four patients were right-and 10 were left-handed. All patients met the following inclusion criteria: 1) First-ever clinical ischemic stroke with a positive DWI lesion within the previous 2 weeks; 2) One-sided upper extremity weakness (MRC < 5); 3) Ability to give informed consent and understand the tasks involved. We excluded patients with one or more of the following criteria: initial UE FMA > 63/66, age under 21 years, hemorrhagic stroke, space-occupying hemorrhagic transformation, bihemispheric stroke, traumatic brain injury, encephalopathy due to major non-stroke medical illness, global inattention, large visual field cut (greater than a quadrantanopia), receptive aphasia (inability to follow 3-step commands), inability to give informed consent, major neurological or psychiatric illness that could confound performance/recovery, or a physical or other neurological condition that would interfere with arm, wrist, or hand function recovery. Due to the exclusion of aphasic patients, the sample had a bias towards right-sided infarcts (17 left-sided, 37 right-sided; for detailed patient characteristics, see Table 1). The lesion distribution is shown in Fig. 5A.

**Table 1.** Patient characteristics: age (years), sex, paretic side, initial FMA (Fugl-Meyer arm score, maximum 66), initial MoCA (Montreal Cognitive Assessment, maximum 30).

| Patient | Age at stroke | Gender | Paretic Side | Initial impairment (FMA) | Initial MoCA |
| --- | --- | --- | --- | --- | --- |
| 1 | 57 | M | R | 48 | 27 |
| 2 | 24 | M | L | 35 | 23 |
| 3 | 67 | F | R | 16 | 23 |
| 4 | 74 | F | R | 39 | 17 |
| 5 | 61 | F | L | 48 | 26 |
| 6 | 59 | F | R | 60 | 28 |
| 7 | 57 | M | R | 54 | 27 |
| 8 | 66 | M | L | 65 | 25 |
| 9 | 42 | F | R | 5 | 18 |
| 10 | 65 | M | L | 30 | 25 |
| 11 | 66 | F | L | 60 | 19 |
| 12 | 51 | M | L | 34 | 25 |
| 13 | 63 | F | L | 57 | 26 |
| 14 | 55 | M | L | 0 | 26 |
| 15 | 56 | M | L | 38 | 25 |
| 16 | 56 | M | L | 64 | 24 |
| 17 | 64 | F | R | 20 | 16 |
| 18 | 60 | F | R | 55 | 21 |
| 19 | 64 | M | L | 63 | 25 |
| 20 | 25 | F | L | 42 | 29 |
| 21 | 39 | F | L | 47 | 20 |
| 22 | 46 | M | L | 9 | 27 |
| 23 | 53 | F | L | 4 | 29 |
| 24 | 66 | M | L | 59 | 24 |
| 25 | 71 | M | L | 4 | 26 |
| 26 | 52 | M | L | 53 | 24 |
| 27 | 46 | M | R | 4 | 21 |
| 28 | 46 | M | L | 49 | 30 |
| 29 | 71 | M | L | 6 | 24 |
| 30 | 47 | M | R | 57 | 10 |
| 31 | 45 | M | L | 8 | 27 |
| 32 | 55 | F | L | 19 | 25 |
| 33 | 68 | F | L | 61 | NaN |
| 34 | 65 | M | L | 32 | 28 |
| 35 | 51 | F | L | 63 | 26 |
| 36 | 42 | M | R | 54 | 25 |
| 37 | 58 | M | L | 4 | 24 |
| 38 | 41 | F | L | 4 | 23 |
| 39 | 35 | M | L | 4 | 29 |
| 40 | 68 | M | L | 52 | 27 |
| 41 | 76 | M | L | 53 | 18 |
| 42 | 86 | M | L | 54 | 20 |
| 43 | 48 | M | L | 16 | 25 |
| 44 | 74 | M | R | 5 | 25 |
| 45 | 80 | F | R | 9 | 24 |
| 46 | 64 | F | L | 58 | 19 |
| 47 | 22 | M | R | 63 | 27 |
| 48 | 88 | F | R | 55 | 28 |
| 49 | 22 | M | R | 63 | 27 |
| 50 | 87 | F | R | 50 | 28 |
| 51 | 84 | M | R | 30 | 26 |
| 52 | 53 | M | R | 30 | 29 |
| 53 | 54 | M | L | 59 | 21 |
| 54 | 58 | M | R | 61 | 23 |

We also recruited 14 age-matched healthy control participants (10 male, 4 female; mean age 64±8.2 years; all right-handed) at the three centers. There was no age difference between the patient and control samples (two-samples t-test *t*(65) = 1.60, *p* = 0.11), nor did the ratio of gender and handedness in the two groups differ (Fisher’s exact test yielded results of p = 0.11 and 0.75, respectively). The healthy controls did not have any neurological disorder or physical deficit involving the upper limbs. All participants signed a written consent, and all procedures were approved by Institutional Research Board at each study center.

### Assessment of finger maximum voluntary contraction and of individuation

To achieve good characterization of hand function recovery, the study design required patient testing at the following five time points post-stroke: within the first 2 weeks (W1, 10±4 days), at 4-6 weeks (W4, 37±8 days), 12-14 weeks (W12, 95±10 days), 24-26 weeks (W24, 187±12 days), and 52-54 weeks (W52, 370±9 days). Healthy controls were tested at comparable intervals.

At each of the five visits, hand function was tested using an ergonomic device that measures isometric forces produced by each finger (Fig. 1A). The hand-shaped keyboard was comprised of ten keys. Force transducers (FSG-15N1A, Honeywell^®^; dynamic range 0-50 N) measured the force exerted by each finger with a sampling rate of 200 Hz. The data were digitized using National Instrument USB-621x devices interfacing with MATLAB (The MathWorks, Inc., Natick, MA) Data Acquisition Toolbox. Visual stimuli of the task were presented on the computer monitor, run by custom-written software using the Psychophysics Toolbox (Psychotoolbox) in MATLAB environment (Brainard, 1997).

**Figure 1.**
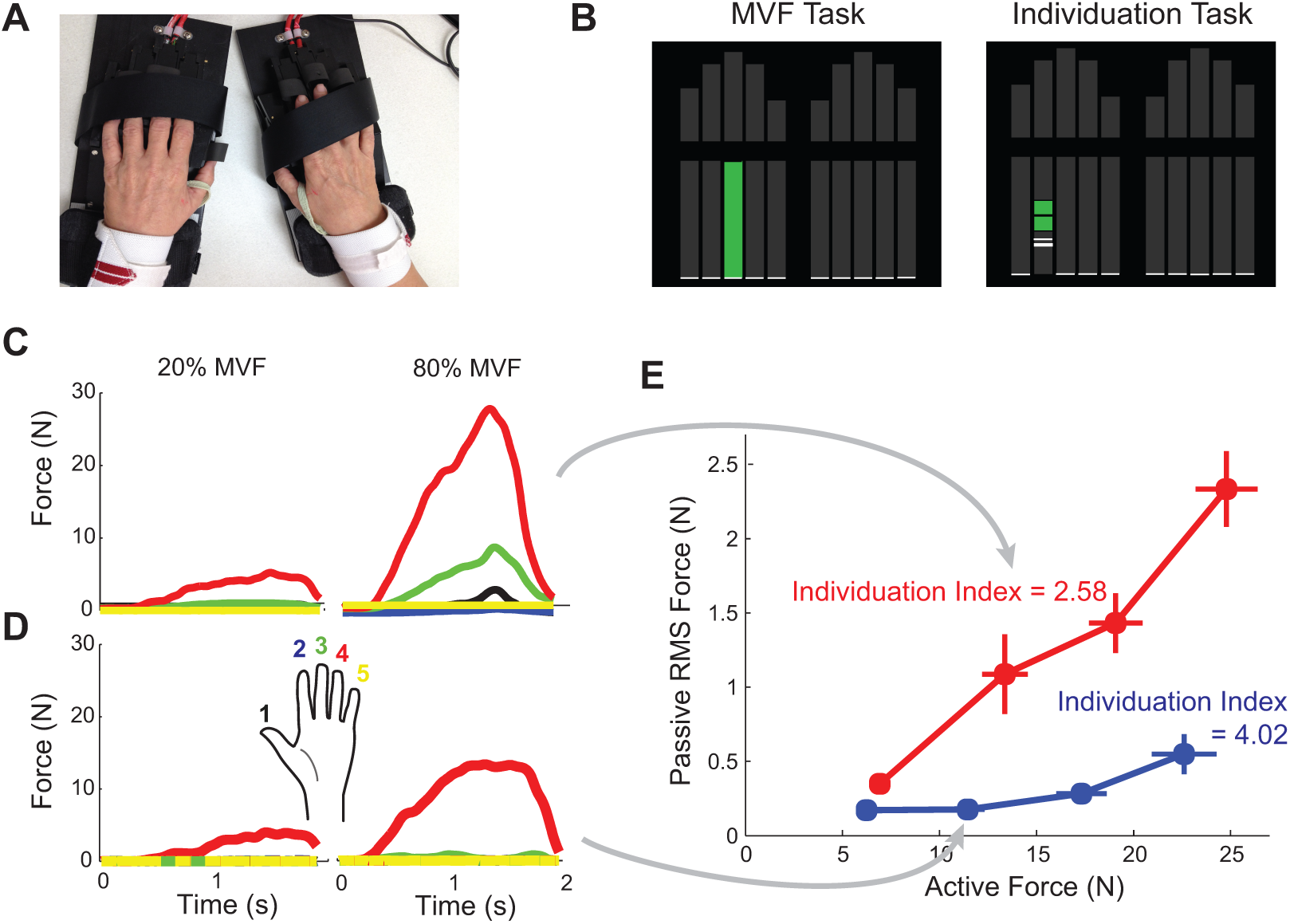
Strength and Individuation task. (**A**) Ergonomic hand device. The participant’s fingers are securely placed on the keys using Velcro straps. (**B**) Computer screen showing the instructional stimulus, which indicates both which finger to press and how much force to produce (height of the green bar). In the MVF task, maximal force was required; in the Individuation task a specific force level had to be reached. (**C**, **D**) Example trials from two healthy control participants during the Individuation task. Four trials are shown, one at 20% and one at 80% of MVF for the two participants. In this case the fourth finger (red) was the active finger. Note the higher level of enslaving of the passive fingers for higher active force level. (**E**) Mean deviation from baseline in the passive fingers plotted against the force generated by the active finger for (C) and (D). Increased enslaving with higher active force levels is clearly visible. The Individuation Index is the -log(slope) of the regression line between active force and passive mean deviation, measured as root mean square (RMS) force from baseline force produced by passive fingers.

Participants were seated in a comfortable chair, facing the computer monitor. During the entire procedure, participants rested their two hands on the keyboards with each finger on top of a key, their wrists were strapped and fixed on a wrist-rest, and their forearms extended and supported by foam arm rests. Throughout the experiment, ten vertical gray bars representing the ten digits appeared on top of the screen, and another ten vertical bars below them instructed the amount of force to be exerted; the required force level for each finger in each trial was indicated by the height of green filling the vertical gray bar (Fig. 1B). Participants could monitor the force exerted by all ten digits in real time by the heights of ten small white horizontal lines moving along the vertical force bars.

Two separate aspects of finger function were tested: maximal voluntary contraction force (MVF) and individuation. During each MVF trial, participants were asked to depress one finger at a time with its maximum strength, and maintain the force level for two seconds. The participants could press with the other fingers as much as they wanted as long as maximal force on the instructed finger was achieved. To signal the start, one force bar corresponding to the instructed finger turned to green. MVF was measured twice per digit.

In the individuation task, participants had to press each individual finger at a sub-MVF level of force, while at the same time keeping their other fingers immobile on the keys. Four target force levels were tested for each digit: 20%, 40%, 60%, and 80% of MVF, and each level was repeated 4 times. On each trial, a section of a force bar corresponding to the finger to be depressed turned to green, with the height of the middle black line representing the target force level and the green region around the middle line representing the 25% upper and lower bounds around the target force level (Fig. 1B). The participants were asked to bring the corresponding white line up to the force target line, and maintain the force level for 0.5 sec. If no response passing the force threshold of 2.5 N was detected within two seconds, the trial was terminated.

### Data analysis

#### Strength Index

The 95th percentile of the force traces produced across all the sampled force data points during the finger depressing period in each trial was calculated, and then averaged across the two MVF trials to obtain a measure of MVF for each digit. If the force achieved on one of the two trials was below 60% of the force produced on the other trial, only the larger force was taken as MVF measure (6.5% of the trials were excluded). The overall strength of the hand was then calculated by averaging across all five digits. To account for the large inter-subject variability in premorbid strength, all MVF values were normalized by MVF of the non-paretic hand at W52; estimated using a mixed-effects model (see below). This normalization provided a Strength Index, with a value close to 1 implying full recovery. For control participants, one of the hands was randomly assigned to take the role as the “non-paretic” hand for normalization purposes. To account for possible laterality effects, the assignment followed the ratio of dominant to non-dominant hands found in the patients (~10:4).

#### Individuation Index

If individuation was perfect, a participant should be able to press the instructed finger without any force being exerted by the passive fingers. For each time bin *t* (5ms) in a single trial, we calculated the enslaved deviation of the force of each passive finger (*F*_*t,j*_) from baseline force (*BF*_*j*_), which was assessed at the beginning of the trial when a go cue was presented. This deviation was averaged over all bins (*T*) in the force trace from the go cue to the end of the trial:

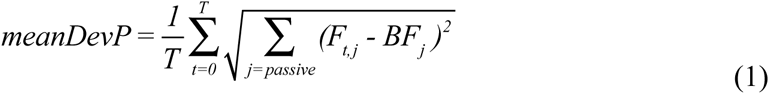

where the index *j* denotes the *j*th passive finger. A higher *mean deviation* indicates more enslaving of the passive finger.

For a measure of individuation ability, it is necessary to account for the relationship between the force deviations of the passive fingers to the force produced by the active finger. Consistent with previous reports (Li *et al.*, 1998), we observed that enslaving of passive fingers increases with higher active force (Fig. 1E). The relationship between the two variables was close to linear. Thus a good measure of individuation is how much the mean deviation in passive fingers increases for each *N* of force produced by the active finger. The ratio of these two variables can be reliably estimated by fitting a regression line without an intercept. To reduce the influence of outliers, we used robust regression (Holland and Welsch, 1977). The *slope* of the regression line reflects individuation ability: The smaller the slope, the better the individuation ability, with the best case being 0, which means keeping the passive fingers perfectly immobile at any active force level. Because the regression slope is bounded by zero (as mean deviation is positive), its distribution is positively skewed. To allow for the use of parametric statistics the slope was log-transformed. The sign of this value was inverted, so that higher values would correspond to better function. The negative log slope was calculated separately for each active finger and then averaged across fingers, giving the Individuation Index for the hand. As was done for the Strength Index, the Individuation Index was normalized by each participant’s “non-paretic” (randomly assigned for healthy controls) hand’s W52 value as estimated by mixed-effect model to provide the final Individuation Index for each hand.

#### Reliability measures for Strength and Individuation

To determine the reliability of the Strength and Individuation Indices, split-half reliabilities for both measures were calculated. For the Strength Index, we used one MVF trial per digit in each split. We then calculated the (normalized) Strength Index on each half of the data independently in the same way as for the full data set. The correlation between the two halves across all available sessions and patients was then used as a measure of split-half reliability.

For the Individuation Index, data from each finger was split such that two trials per force level were assigned to each split. The slope of the regression line and Individuation Index was then calculated separately for each split, and normalized in the same way as MVF. We repeated the split multiple times, each time assigning trials at random and then averaging the split-half correlations from all splits for more reliable results.

Split-half correlation will underestimate reliability because the variability in each half will be higher than the variability when using all the data (Guttman, 1945). The estimate was therefore corrected using the formula

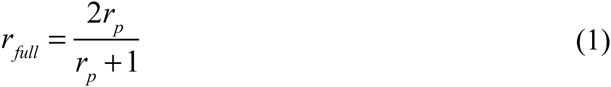

where *r*_*p*_ is the correlation between the two splits.

#### Stability analysis

To assess whether the relative deficits in strength and individuation remained stable across different testing time points, or whether there was meaningful biological change, we calculated the correlation of each measure across neighboring testing time points. One caution when interpreting these correlations is that the correlation between two repeated measures will always be smaller than 1 even if the underlying factor did not change. This is because both measures contain some measurement noise. To account for this effect, we used the reliability (*r*_*full*_) of the measure at each time point to compute a noise ceiling, which indicates how much two repeated noisy measurements should correlate with each other if the underlying variable were perfectly stable:

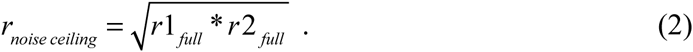

#### Statistical analysis and handling of missing data

Data analysis was performed using custom-written MATLAB and R (R Core Team, 2012) routines. The analysis focused on the Strength and Individuation Indices, but was also performed on standard clinical assessments, FMA, ARAT, and Dynamometry strength measures.

The requirement for 5 post-stroke time-points was ambitious, with the consequence of some missing sessions. A total of 21 patients completed all five time-points; on average each patient completed 3.6 sessions; thus a total of 75% of the possible sessions were acquired. To optimally use all the measured data, we employed linear mixed-effect models. The model specifies joint distributions for observed and missing observations – then the parameters of those distributions can be estimated by maximizing the likelihood of the data under the model. There are several advantages to this approach. First, all the available data can be used and there is no need to exclude any data.

Secondly, it avoids the statistical pitfalls inherent in “filling in” missing observations with point estimates. Linear mixed-effect models implemented in the *lme4* package in R (Bates *et al.*, 2014) were used to test the changes in these measures over time. Participant was taken as a random factor. Time Point (five time points from W1-W52) and Hand Condition (paretic, non-paretic, and control) were considered fixed factor. The model was applied to control and patient data separately. Mixed-effect model estimation for group summary statistics was implemented in MATLAB using the restricted maximum likelihood method (Laird and Ware, 1982).

#### Modeling the time-invariant function

To test the hypothesis that there is time-invariant relationship between strength and individuation, a two-segment piecewise linear function was fitted. This function had four free parameters: the intercept, the location of the inflection point, and the slope on each side of the inflection point. Let *x* be the predictor with two segments separated by a constant breakpoint *c*, *x*_1_ ≤ *c* and *x*_2_ ≥ *c*. The linear functions for each segment are

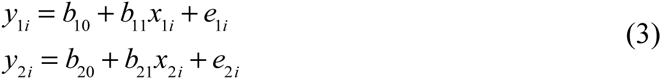

The two pieces can be joined at the breakpoint constant c by setting *y*_*1i*_ = *y*_*2i*_, yielding

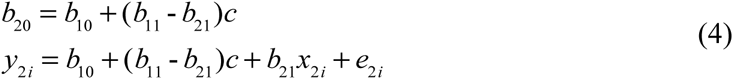

Putting the two pieces together, we have the full model

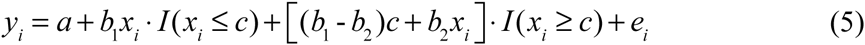

where *I* (⋅) is an indicator variable, coded as 1’s or 0’s to indicate the condition satisfied.

The maximum-likelihood (or least-squares) estimates of these parameters were obtained by using the non-linear optimization routine fminsearch in Matlab. Leave-one-out cross-validation (Picard and Cook, 1984) was used to evaluate whether this function changed systematically over time, or whether it was time-invariant. The time-invariant model with fixed parameters across all time points was compared with a more complex model that allowed free parameters for each time point. Cross validation provides an unbiased estimate of a model’s ability to predict new data and automatically penalizes models that are too complex.

### Lesion Imaging and Quantification

#### Imaging acquisition and lesion distribution

Images were acquired using 3T MRI Phillips scanner and consisted of two DTI datasets (TR/TE=6600/70ms, EPI, 32 gradient directions, b=700 s/mm2), and an MPRAGE T1-WI (TR/TE=8/3.8ms) sequence. FOV, matrix, number of slices, and slice thickness were 212×212 mm, 96×96 (zero-filled to 256×256), 60, 2.2mm, respectively, for DTI; and 256×256mm, 256×256, 170, 1.2mm, respectively, for T1-WI. The DTI were processed using DtiStudio (www.MRIStudio.org) and the mean diffusion-weighted image (DWI) was calculated.

To define the boundary(s) of the acute stroke lesion(s) for each participant, a threshold of >30% intensity increase from the unaffected area in the first-time-point diffusion-weighted image (DWI) extracted from DTI images was applied. A neuroradiologist (AVF), blind to the patients’ clinical information, manually modified the boundary to avoid false-positive and false-negative areas on RoiEditor (www.MRIstudio.org). The definitions were double checked by a second rater (MB). The averaged lesion distribution map across all patients in the current study is shown in Fig. 5A. For the seven patients who had no DTI in the acute phase, lesion definition was performed on the clinical DWI, which has lower resolution (1×1mm in plane, 4-6mm thickness). Analysis of white matter ROIs, including the CST, was not performed in patients.

#### Region of interest definition and lesion quantification

The focus was on two ROIs: 1) The cortical gray matter of the hand area in the motor cortex; 2) The entire CST superior to pyramids, identified by probabilistic maps derived from tract tracing methods (see below). The percentage volume affected in these regions was correlated with our main outcome measures, the Strength and Individuation Indices.

To defined the CST, each image and respective lesion were mapped to a single subject adult template, the JHU-MNI atlas (Mori *et al.*, 2008; Oishi *et al.*, 2008, 2009, 2010), using affine transformation followed by dual channel (both b0 and FA maps) large deformation diffeomorphic metric mapping (LDDMM) (Ceritoglu *et al.*, 2009). This template has already been segmented into more than 200 regions of interest (ROIs), and contains probabilistic maps of multiple tracts, including the CST (Zhang *et al.*, 2010). To ensure accurate mapping, we first used "artificial" images, in which the stroke area was masked out and substituted by the normal images from the contralateral hemisphere. This helped to minimize inaccuracies caused by the focal changes in intensity due to the stroke. The white matter beneath the cortex was identified with a FA-threshold of 0.25. The segmentation defined in the template, as well as a probabilistic map of the CST, were then “back-warped” by each subject's deformation field to generate individualized parcellations.

A different approach was used to define an ROI that would encompass the hand area of the primary motor cortex. The hand ROI was defined on the average reconstruction of the cortical surface available in the Freesurfer software (Dale *et al.*, 1999), selecting Brodmann area (BA) 4 based on cytoarchitectonic maps (Fischl *et al.*, 2008). To restrict the ROI to the area of motor cortex involved in the control of the upper limb, we only included the area 2.5 cm dorsal and ventral of the hand knob (Yousry *et al.*, 1997). The defined ROI was then morphed into MNI space using the surfaces of the age-matched controls. These ROIs were then brought to the JHU-MNI atlas (in which each subject and respective stroke area were already mapped, as mentioned above) using T1-based LDDMM to construct a probabilistic map of the hand area. The probabilistic map was threshold of 70% to calculate percent-volume affected.

### Clinical assessments

At each visit, all participants were also assessed with several clinical outcome measures. Here we report data for FMA and ARAT. Grip strength was assessed with a Jamar hydraulic hand dynamometer (Sammons Preston, Rolyan, Bolingbrook, IL, USA). Strength in the first dorsal interosseous (FDI) and the flexor carpi radialis (FCR) muscles was assessed using a hand-held dynamometer (Hoggan MiroFET2 Muscle Tester, Model 7477, Pro Med Products, Atlanta, GA, USA).

## Results

A total of 54 patients with acute stroke and 14 healthy controls underwent five testing sessions over a one-year period. Data in the final analysis comprised a total of 251 sessions tested in 53 patients (one patient only completed two blocks of the task, and was thus removed from further analysis) and 14 controls. Forty-one patients and twelve controls completed >= 3 sessions. The data were 75% complete, with 25% of the sessions were missing or unusable. Non-tested sessions were treated as data missing at random and all available data were used in the statistical analysis (see Materials and Methods).

### The Strength and Individuation Indices were reliable

Finger strength was assessed by measuring the maximum voluntary force (MVF) for each finger separately and then averaged across all fingers for each hand. MVF for healthy controls had an average value of 20.35 N (SD = 8.56) for the dominant hand, and 22.76 N (SD = 6.89) for the non-dominant hand. The normalized Strength Index for the controls’ dominant hand was 1.00 (SD = 0.19), and non-dominant hand was 1.17 (SD = 0.25). For patients, the mean for the non-paretic hand was 0.93 (SD = 0.20), and for the paretic hand it was 0.59 (SD = 0.38). For the paretic hand, Strength Indices did not correlate with age (*r* = 0.04, *p* = 0.75), nor were they affected by gender (*t*(51) = 0.98, *p* = 0.33) or handedness (*t*(51) = 0.10, *p* = 0.92).

To assess individuation, we measured the amount of involuntary force changes (enslaving) on the passive fingers for different levels of force production with the active fingers. The amount of enslaving systematically increased at higher force levels (Fig. 1E). Loss of control at increasing force levels has been shown for the angular position of the fingers (Li *et al.*, 1998) and the reaching radius of the arm after stroke (Sukal *et al.*, 2007; Ellis *et al.*, 2009). To control for this relationship, we characterized the Individuation Index as the slope of the function between active force and passive enslaving. Lower values of Individuation Index indicate more impaired individuation. Healthy, age-matched controls showed, on average, a normalized Individuation Index of 1.00 (SD = 0.18). This refers to a slope of 0.087 (SD = 0.046), meaning that for a finger press of 10N the mean deviation of the passive fingers was 0.69N. As was the case for Strength, Individuation Indices in the paretic hand were not correlated with age (*r* = 0.16, *p* = 0.26), nor affected by gender (*t*(51) = 0.17, *p* = 0.86) or handedness (*t*(51) = 0.34, *p* = 0.74).

When introducing a new instrument, it is important to first establish its reliability, i.e., the accuracy with which true differences between subjects and changes within subject can be determined. We therefore split the data for each session in half, calculated Strength and Individuation Indices on these two independent data sets, and correlated the resultant scores across patients and sessions (see methods). The adjusted split-half reliability across all patients and weeks for the Strength Index was *r*_*full*_ = 0.99 and 0.94 for the paretic and non-paretic hands respectively, and *r*_*full*_ = 0.89 for controls, which indicates good reliability. The adjusted split-half reliability of the Individuation Index of all patients was *r*_*full*_ = 0.99 and 0.93 for the paretic and non-paretic hands respectively, and for controls was *r*_*full*_ = 0.97.

Consistent with our effort to construct an individuation measure that is independent of strength, the overall correlation between Individuation and Strength in controls was very low for both controls (*r* = -0.19, *p* = 0.51), and for patients’ non-paretic hand (*r* = 0.17, *p* = 0.21).

### The Strength and Individuation indices correlated with standard clinical measures

The Strength and Individuation Indices were compared with existing clinical measures: the Fugl-Meyer (a measure of impairment) and ARAT (a measure of activity) Table 2 shows the correlations for all four measures obtained from the paretic hand across all time points. Overall, all correlations were very high (max *p* = 1.21×10^−26^), indicating that all the measures could detect severity of the hand function deficit. The correlation in the patients between the two clinical measures was 0.91, whereas the correlation between the Strength and Individuation Indices was 0.73, a significant difference (*z* = 5.62, *p* = 2.0×10^−8^, using *z*-test with N = 180 (Fisher, 1921)). Given comparable reliabilities for all measures, this difference unlikely results from measurement noise – rather it suggests that our Strength and Individuation Indices measure two different aspects of the hand function, whereas the clinical scales tend to capture a mixture of strength and control.

**Table 2.** Correlation between Strength Index, Individuation Index, FMA (Fugl-Meyer arm score, maximum 66), and ARAT (Action Reach Arm Test, maximum 57). All four measures are highly correlated; however Strength and Individuation Indices show the weaker correlation compared to that between FMA and ARAT.

|  | Strength<br>Index | Individuation<br>Index | FMA | ARAT |
| --- | --- | --- | --- | --- |
| Strength<br>Index |  | 0.73 | 0.76 | 0.74 |
| Individuation<br>Index |  |  | 0.68 | 0.72 |
| FMA |  |  |  | 0.91 |

### Recovery of strength and individuation occurred mainly in the first three months after stroke

We first examined the time courses of recovery for strength and individuation in the paretic hand. If the two observed variables change in parallel, their recovery may or may not be mediated by the same underlying process. A difference in the time courses, however, would provide a strong hint of separate recovery processes for strength and individuation.

For both measures, most of the recovery appeared to occur within the first 12 weeks after stroke (Fig. 2A-B). A model with a fixed effect of Week and a random effect of Subject was built to evaluate this statistically. An effect of Week was tested with a likelihood ratio test against the null model with the random effect only. Results indicate that both the Strength and Individuation Indices significantly improved over time (Strength: *χ*^2^ = 47.65, *p* = 5.10×10^−12^; Individuation: *χ*^2^ = 18.58, *p* = 1.63×10^−5^). Paired *t*-tests between adjacent time points showed significant improvement (after Bonferroni correction) of the Strength Index up to week 12; whereas the Individuation Index only showed a significant improvement between weeks 4-12 (see detailed statistics in Fig. 2A-B). A similar recovery curve was found for the standard clinical measures of motor function (detailed statistics in Fig. 3).

**Figure 2.**
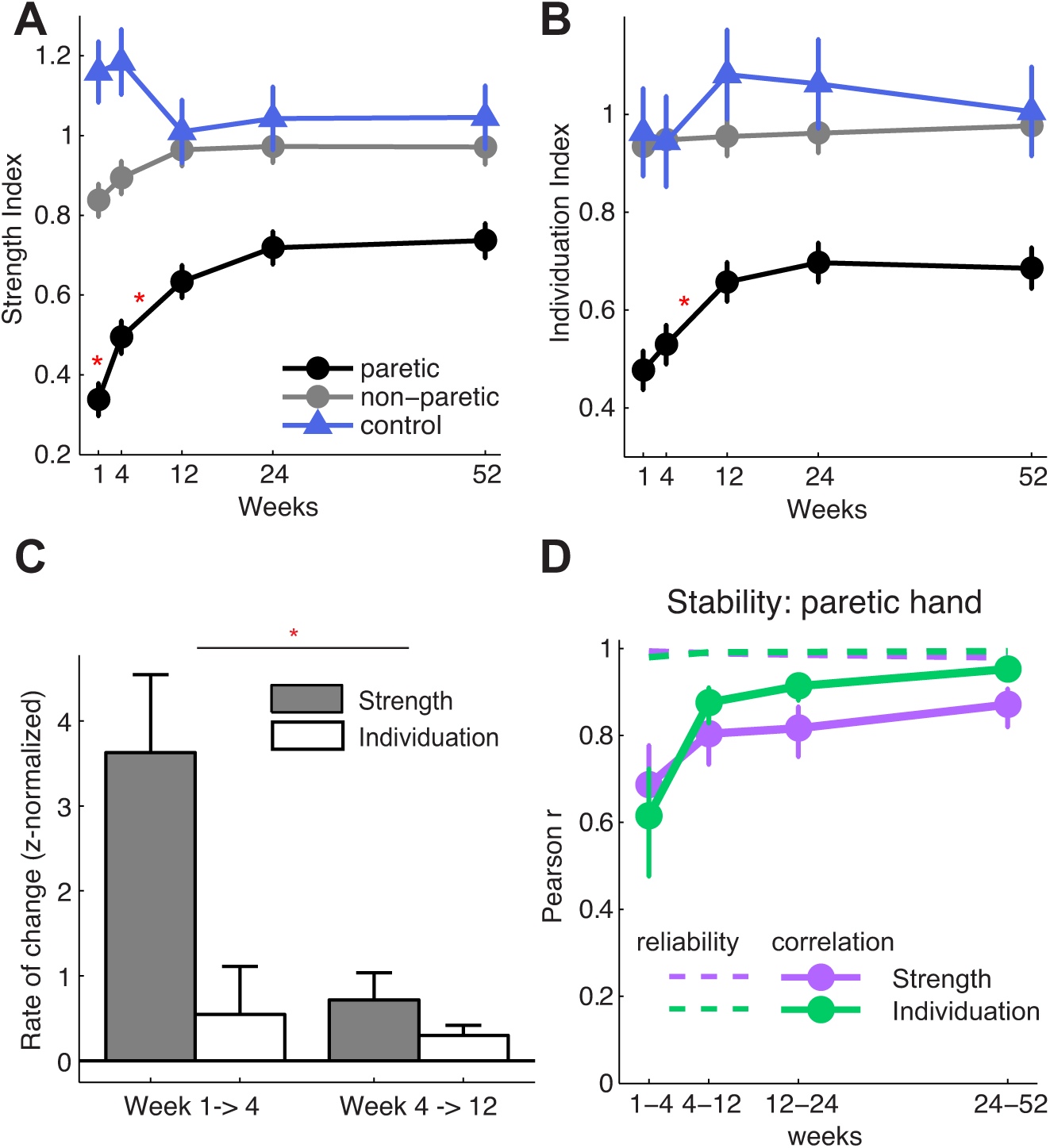
Temporal profiles of recovery for strength and individuation. (**A-B**) Group recovery curves for the Strength and Individuation Indices for patients and controls. Asterisks indicate significant week-to-week change for the paretic hand (Bonferroni corrected p-values for each segments of Strength Index: p(W1-4) = 0.0045, p(W4-12) = 0.0082, p(W12-24) = 0.068, p(W24-52) = 0.87; Individuation Index: : p(W1-4) = 0.81, p(W4-12) = 0.0024, p(W12-24) = 1.92, p(W24-52) = 2.91). (**C**) Rate of change (i.e., change per week) in Z-normalized Strength and Individuation Indices during the first two intervals (Week 1 to 4 and Week 4 to 12). The two intervals show a significant interaction between strength and individuation, indicating faster initial improvement of strength; (**D**) Week-to-week correlations between adjacent time points for the Strength and Individuation Indices. Dashed lines are the noise ceilings based on the within-session split-half reliabilities.

**Figure 3.**
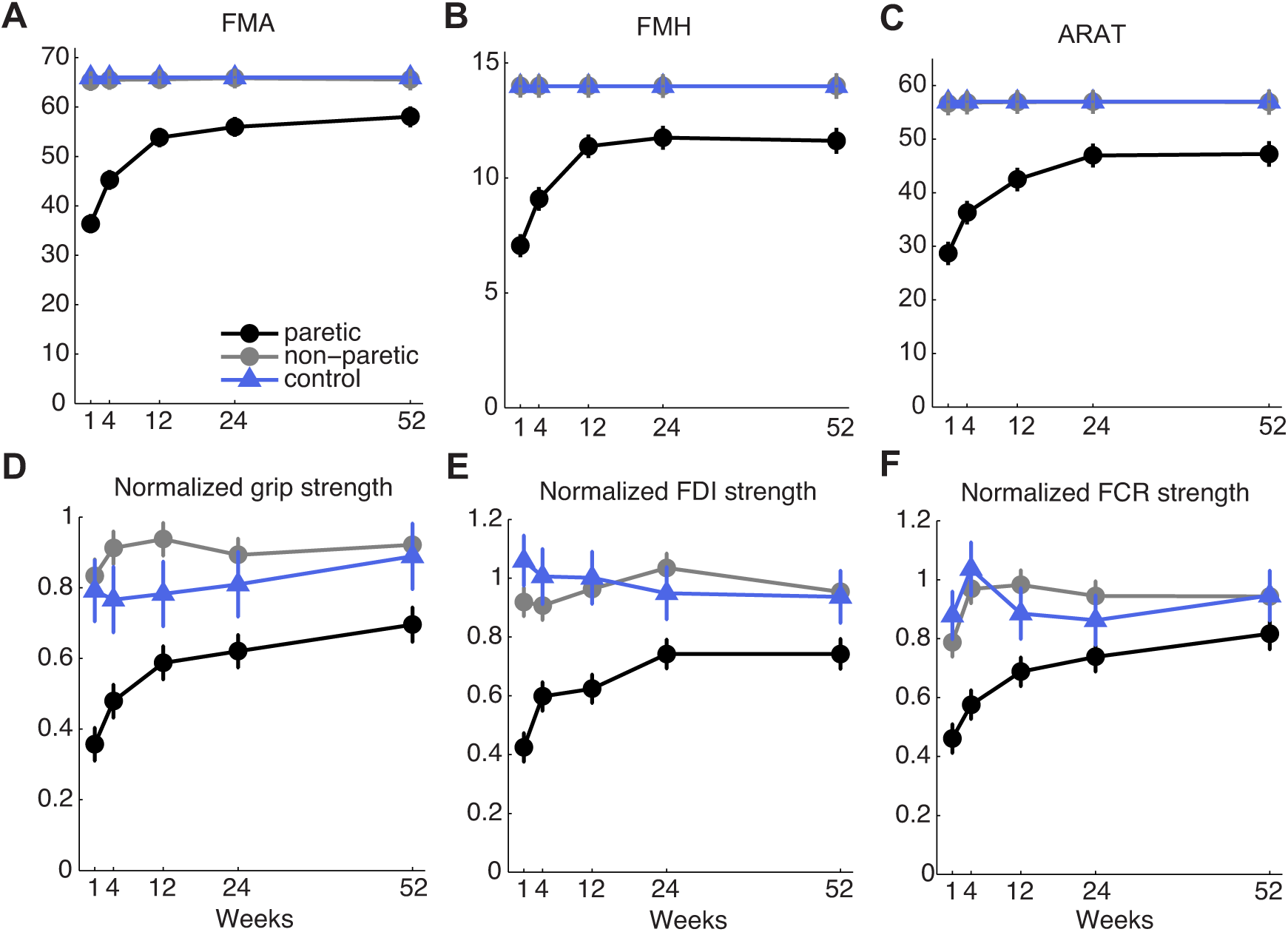
Temporal recovery profiles measures with clinical assessments. (A) Fugl-Meyer for the arm (FMA) and (B) hand (FMH); (C) ARAT; (D-F) strength for hand grip, FDI, and FCR muscles, as measured by Dynamometry. All measures showed significant change over time for the paretic hand. *FMA*: χ^2^ = 37.73, *p* = 8.13×10^−10^; *FMH:* χ^2^ = 29.03, *p* = 7.14×10^−8^; *ARAT:* χ^2^ = 36.33, *p* = 1.67×10^−9^; *grip:* χ^2^ = 33.02, *p* = 9.21×10^−9^; *FDI:* χ^2^ = 19.21, *p* = 1.67×10^−5^; *FCR:* χ^2^ = 28.47, *p* = 9.50×10^−8^.

To directly compare the time courses of between the two indices at the early stage of recovery, we *z*-normalized scores of the two variables and then investigated the change in the scores for the time intervals W1-4 vs. W4-12 (Fig. 2C). This analysis suggests that strength may recover mostly in the first four weeks, while individuation recovery may occur equally in both time periods. Repeated-measures ANOVA over z-scores for Strength and Individuation Indices during the two time intervals yielded a significant interaction (*F*(1,25) = 6.82, *p* = 0.015, Fig. 2C). Thus, despite overall similarity, there was a significant difference in the time courses of recovery of strength and individuation, with strength showing faster early recovery.

That most improvement in both strength and individuation occurred over the first 12 weeks is also apparent in the correlations between adjacent testing time points for each variable across individuals (Fig. 2D). The correlation between weeks 1 and 4 for the Individuation Index was significantly lower than it was for subsequent time points (W1-4 vs. W24-52: *z* = -4.23, *p* = 0.000023), and this difference for Strength Index was marginally significant (*z* = -1.83, *p* = 0.067), using *z*-test with N = 28 and 33 (Fisher, 1921). Thus, the position of the patients on the mean recovery curve changed more during the first 4 weeks than in the last 6 months. This correlation difference cannot be attributed to measurement noise, as both measures had stable reliabilities at all time points (dashed line). Instead, the lack of stability of these measures during early recovery is indicative of meaningful biological change.

Consistent with previous findings (Noskin *et al.*, 2008), the non-paretic hand also showed mild impairment in the first month after stroke. A likelihood ratio test of the mixed-effect model showed a significant effect of Week for Strength (*χ*^2^ = 7.86, *p* = 0.0051), and a more subtle effect for Individuation (*χ*^2^ = 4.12, *p* = 0.042) (Fig. 2A-B). This increase in performance is unlikely to be related to a general practice effect, because the Strength Index in healthy controls decreased slightly over time (*χ*^2^ = 4.54, *p* = 0.033), perhaps due to reduced effort, whereas the Individuation Index for healthy controls was maintained at a similar level over the whole year (*χ*^2^ = 0.33, *p* = 0.56).

In summary, most recovery of both strength and individuation occurred in the first three months after stroke, with stabilization of recovery around 3-6 months. The data also suggest a slight difference in the time course, with strength recovering faster than individuation in the first month.

### Evidence for a time-invariant relationship between strength and control

The time course analysis only provides weak evidence for partial independence of the recovery processes for strength and individuation. Therefore we undertook a closer examination of the relationship between the two variables at each testing time-point (Fig. 4A). Although patients tended to move from the lower left corner to the upper right corner of this space over the time course of recovery, the overall shape of the strength-individuation impairment relationship seemed to be remarkably preserved across weeks. At lower strength levels, there was a clear correlation between strength and individuation; whereas once above ~60% of normal strength level, the two variables were unrelated, producing a distinct curvilinear shape for the overall function (Fig. 4B).

**Figure 4.**
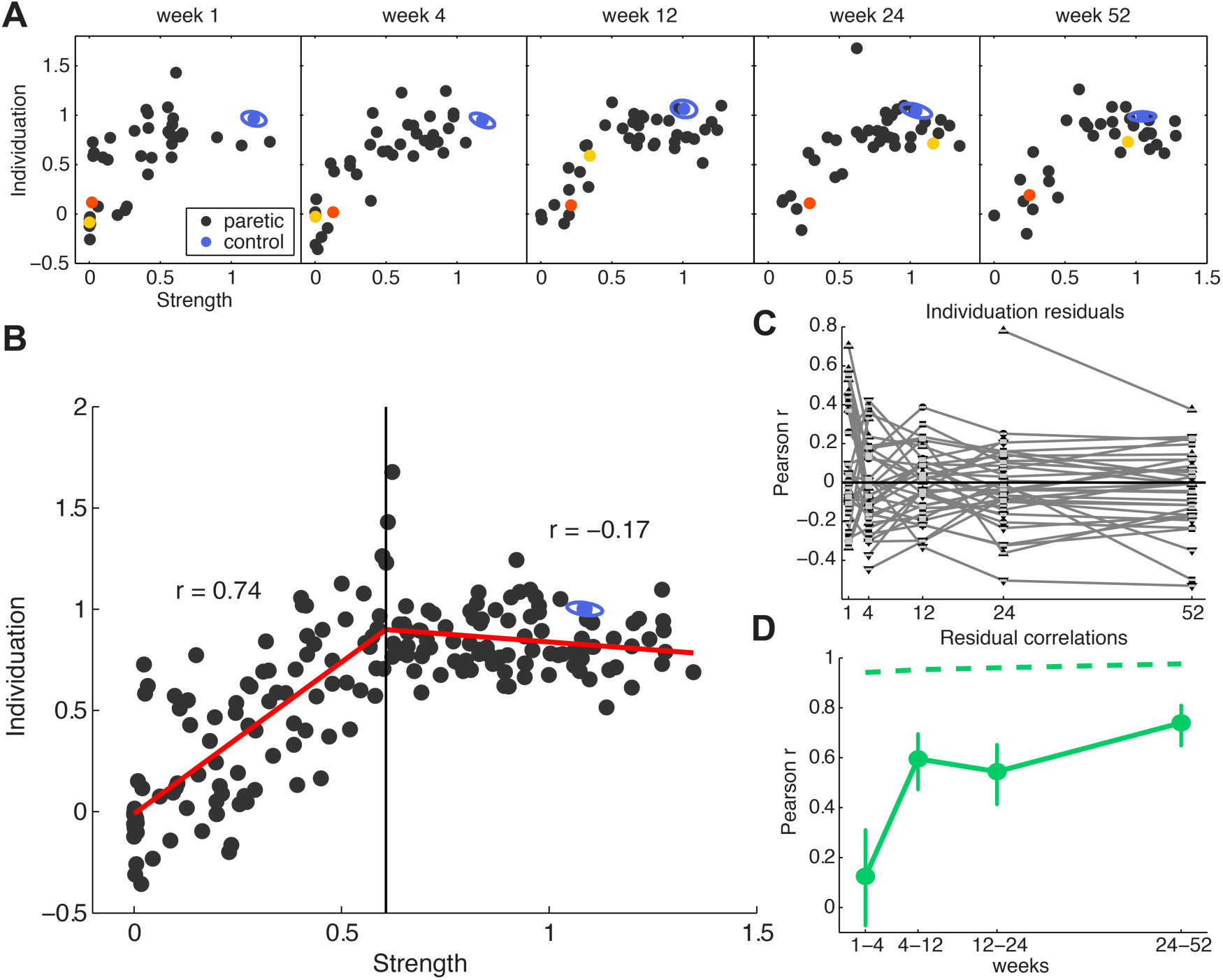
Time-invariant impairment function relating strength and control. (**A**) Scatter plots for Individuation against Strength Indices at each time point. Each black dot is one patient’s data; blue dots and ellipse indicates the mean and standard error for controls at the time point. Two patients’ data are highlighted: one with good recovery (yellow dot) and one with poor recovery (red dot). (**B**) Scatter plot with data from all time points superimposed with the best fitting two-segment piecewise-linear function with one inflection point at Strength Index = 0.607. (**C**) Residuals from each week subtracting out the mean impairment function (B, red line). The tendency of the residuals to stay above or below the typical Strength-Individuation relationship indicates that there are stable factors that determine Individuation recovery over and above strength recovery. (**D**) Correlations of residuals from (C) across adjacent time points increased over time (Bonferroni corrected p(W1-4) = 2.12, p(W4-12) = 0.00064, p(W12-24) = 0.0024, p(W24-52) = *p* = 3.39×10^−6^). Dashed line is the noise ceilings based on the within-session split-half reliabilities.

To formally test the time invariance suggested by visual inspection, we first found a function to describe the strength-individuation relationship. We used data from all time points and evaluated the goodness of fit of a piecewise function with two linear segments connected at an inflection point, using leave-one-out cross-validation (see Materials and Methods). Cross-validation automatically penalizes models that are too complex. This functional form gave us a good fit to the data (cross-validated *R*^*2*^ = 0.53, Fig. 4B). We also explored first- to fourth- order polynomial functions. All four models resulted in a worse fit (cross-validated *R*^*2*^ < 0.49) than the piece-wise linear function.

We then tested for “time-invariance” of this strength-control relationship, that is, whether the function shape changed across weeks. Again, using leave-one-out cross-validation, the time-invariant model with fixed parameters across all weeks was compared with a model that allowed the parameters to change for each week (time-varying model). The cross-validated *R*^*2*^ for the time-varying two-segment piecewise linear function was 0.45, a worse fit than the time-invariant model.

These results suggest that there is a time-invariant recovery relationship between strength and individuation after stroke, which consists of two parts: up to a certain level of strength (60.7% of non-paretic hand), the Strength and Individuation Indices are strongly correlated (*r* = 0.74, *p* = 6.61× 10^−18^); after strength exceeds this threshold, the two variables are no longer correlated (*r* = -0.17, *p* = 0.11; Fig 4B). This lack of correlation cannot be attributed to a ceiling effect for the Individuation Index, because for both patients and controls there was still a considerable variability, and the reliability of Individuation Index was very high. This indicates that our measure has enough dynamic range and sensitivity to detect inter-individual differences even in the healthy population.

Overall, our results suggest that recovery can be captured as traversal along a time-invariant function relating strength and individuation. Differences in recovery arise because patients vary substantially in the distance they move along this function: some patients with initial severe impairment made a good recovery, moving past the inflection point of 60.7% strength (exemplified by the yellow dot in Fig. 4A). Other severely impaired patients failed to reach the inflection point (red dot in Fig. 4A). Finally, some mildly impaired patients started off beyond the inflection point and showed a good range of individuation capacity.

### A second process contributed additional recovery of finger individuation

The fact that recovery of both strength and individuation could be captured by a single time-invariant function that relates them is compatible with the hypothesis of a single underlying process that drives recovery of both aspects of hand function. It is possible, however, that an additional process injects further recovery, which determines a patient’s position relative to the mean recovery function in strength-individuation space. If such a process exists, a given patient should occupy a consistent position above or below the mean recovery function across time points.

To test this hypothesis we investigated the residuals of the Individuation Index for each patient at each time point after subtracting out the mean two-segment piecewise-linear recovery function. If the variability around this mean function were purely due to noise, we should observe no consistent week-by-week correlation between residuals for each patient. Alternatively, if the residuals were to be correlated across weeks, it would indicate that some patients were consistently better at individuation than that predicted from the function, and others were consistently worse, suggesting an additional factor mediating individuation recovery (black arrows in Fig. 6).

Correlations of residuals from adjacent time points across patients were initially quite low. However, from week 4 onwards, most patients’ distances from the mean function remained stable (Fig. 4C-D). This consistent structure in residuals provides evidence for an extra factor contributing to recovery of individuation. The consistent pattern of residuals at later time points could not be attributed to pre-morbid inter-individual differences, because both the Strength and Individuation Indices were normalized to the non-paretic hand. The low week-by-week correlations between early time points argues that the later correlations do not simply reflect sparing of a particular neural system after the stroke. If this had been true, the correlation between the Individuation residuals should have remained constant across all time points. Furthermore, the lower early correlation cannot be attributed to measurement noise, as reliabilities for the early measurement points were high (Fig. 4D). Rather, the initially low but then increasing correlation indicates an additional recovery process operating above the lower bound of the strength-individuation function (Fig. 6). This process is mostly active in the first three months after stroke and determines how well individuation recovers above that expected from the time-invariant recovery function.

### Lesions involving motor cortex and the corticospinal tract correlated more with individuation than strength

To investigate the underlying neural substrates of recovery processes, we correlated the location and size of the lesion with the Strength and Individuation Indices. We were especially interested in the particular role of the corticospinal tract (CST).

While both corticospinal and corticoreticular projections originate in part from the precentral gyrus and are intermingled to some degree, cortical projections to the reticular formation have a more widespread bilateral origin from other pre-motor areas (Keizer and Kuypers, 1989), whereas direct corticospinal projections to ventral horn neurons primarily arise from the anterior bank of the precentral gyrus/central sulcus, i.e. “new M1” (Rathelot and Strick, 2009; Witham *et al.*, 2016). We therefore predicted that extent of the damage to the hand area of the primary motor cortex, and to the white matter ROI that characterizes the most likely course of the CST (see Materials and Methods) would correlate more with Individuation, and less with Strength. Furthermore, lesions in these areas should correlate with individuation recovery over and above the level expected from the mean recovery function.

As hypothesized, the extent of involvement by the lesion of the cortical hand area correlated significantly with the Individuation Index at all time points. For the CST, all correlations were significant after week 1 (Fig. 5B-D). While both lesion measures also correlated with the Strength Index, these correlations were weaker (repeated-measures ANOVA showed a significant main effect for behavioral measure (*F*(1,3) = 146, *p* = 0.001). This difference was not due to measurement noise, as the Strength and Individuation Indices had comparable reliabilities. Furthermore, percent lesion involvement also significantly correlated with the Individuation Index, after accounting for the average Strength-Individuation relationship (*p* < 0.05 for correlations after week 24 for cortical hand area, and after week 12 for CST). Indeed, at W52, the correlations with the residuals were as high as with the Individuation Indices themselves (*r* = 0.61 vs. *r* = 0.57 for the cortical hand area, *r* = 0.51 vs. *r* = 0.54 for the CST). Together these results suggest that Individuation recovery is most heavily determined by the sparing in the hand area of the primary motor cortex and of direct CST projections, while strength recovery may also depend on other spared descending pathways.

**Figure 5.**
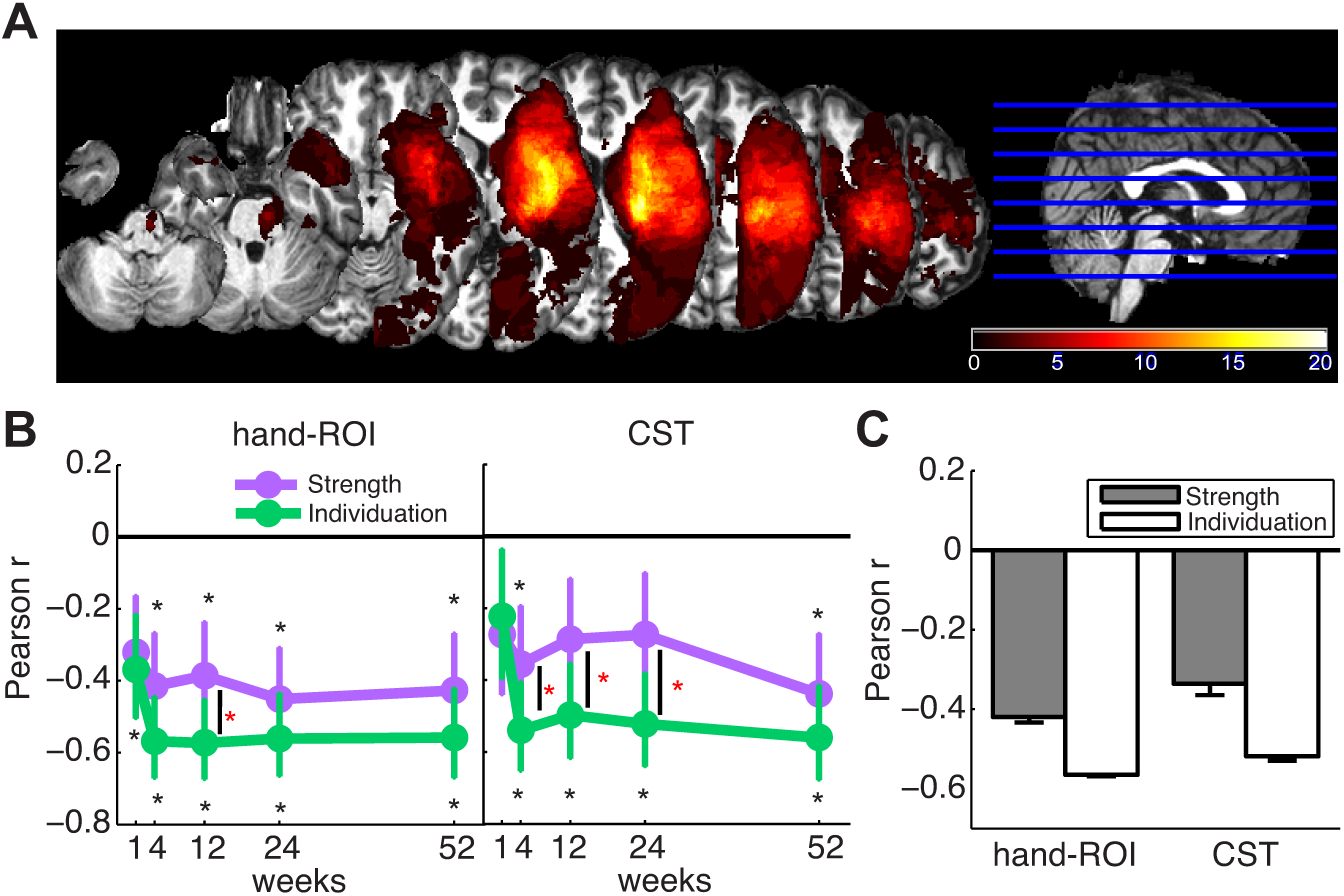
Lesion distribution and correlation with behavior. (**A**) Averaged lesion distribution mapped to JHU-MNI space (see Materials and Methods), with lesion flipped to one hemisphere. Color bar indicates patient count. **(B)** Correlation of behavior measures (Strength and Individuation Indices) at each time point with the percentage of damaged cortical gray matter within the M1 hand area ROI, corticospinal tract (CST). (D) Mean of week-by-week correlations between the two behavior measures and percent lesion volume measures for the cortical gray matter hand area and CST ROI. Black asterisks indicate significant correlations (tested against zero), and red asterisks indicate a significant difference between the correlation for Strength and Individuation for each week (p<0.005).

## Discussion

In a large-scale longitudinal study, we tracked recovery of two independent behavioral components of hand function: strength and finger control. Patients were tested at five time points over a one-year period after stroke, using a novel paradigm that separately measures maximum voluntary contraction force (a measure of strength) and finger individuation ability (a measure of control), and crucially controls for any obligatory dependency between these two measures. This approach allowed us to determine how recovery of strength and control interrelate. Our main question was to ask whether there is a causal relationship between strength and control at the level of recovery mechanisms, after the two variables had been experimentally uncoupled. If they are truly dissociable, then hypothetically patients could show perfect control of individual fingers, even with significant weakness (except for complete hemiplegia, in which case no individuation measure would be obtainable).

We showed that involuntary movements in passive fingers (enslaving) increased with the level of force production of the active finger. This phenomenon is analogous to what Dewald and colleagues (Sukal *et al.*, 2007; Ellis *et al.*, 2009) have described for the paretic arm: a decrease in arm reaching workspace as the force requirement to resist gravity increases. We showed that the ratio of enslaving and active force can account for this dependency and thereby provides a sensitive measure of finger control independent of the level of force deficit.

We first examined the time courses of recovery for strength and individuation. Consistent with what has been described with traditional clinical scales (Duncan *et al.*, 1992; Jørgensen *et al.*, 1995; Krakauer *et al.*, 2012), both measures showed most recovery over the first three months after stroke. This similarity between the time courses does not, however, necessarily imply that recovery of strength and individuation is dependent on a single underlying neural substrate or mechanism. It remains possible that recovery of these two components occurs in parallel because of commonalities in basic tissue repair mechanisms post-ischemia but they are nevertheless independent modules. Indeed, we found a small but robust difference in the time course of recovery of strength compared to control: finger strength showed a faster rate of change compared to individuation over the first month. This finding raises the interesting possibility that different neurological substrates underlie recovery of strength and individuation.

Closer examination of the two variables revealed a time-invariant non-linear relationship between strength and individuation in the paretic hand. This function has two distinct parts: individuation and strength were highly correlated below a strength threshold of ~60% of the non-paretic side; beyond this point, they were uncorrelated. The shape of this function remained the same across all time points. Recovery of hand function could be characterized as movement along this invariant function: patients with good recovery traveled further along the function, whereas patients with poor recovery remained in the first segment. The strong correlation between strength and individuation for severely impaired patients is consistent with a single system mediating recovery of both. Indeed, in our cohort there was no patient with relatively good strength but severe impairment of individuation, which also suggests that recovery of finger control correlates with recovery of strength in patients with severe hemiparesis. However, two pieces of behavioral evidence suggest that strength and finger control might rely on partially separate mechanisms of recovery. First, a correlation between strength and individuation was absent for the subset of well-recovered patients – i.e. patients with a Strength Index above 60%. This breakdown in correlation cannot be attributed to a ceiling effect for Individuation. Secondly, analysis of the residuals around the mean recovery function revealed that patients differed consistently in the amount of their individuation recovery relative to the level predicted by their strength recovery. Notably, their positioning relative to the mean recovery curve seemed to be set early in the recovery process and then remained relatively stable at later time points.

Thus we propose that recovery of strength and individuation relies on at least partially separate systems. One system primarily contributes strength, but also has some limited control capacity. The isolated contribution of this system would determine the lower bound of the data points in the strength-individuation plot (dashed line in Fig. 6): when a patient regains some strength, he or she automatically regains a limited amount of control with it. However, the amount of individuation is limited and does not increase above a certain level. This would explain both the strong correlation between strength and individuation for the severely impaired patients, and the fact that no patient occupied the lower right corner of strength-individuation space, i.e. no patients had good strength but minimal control.

**Figure 6.**
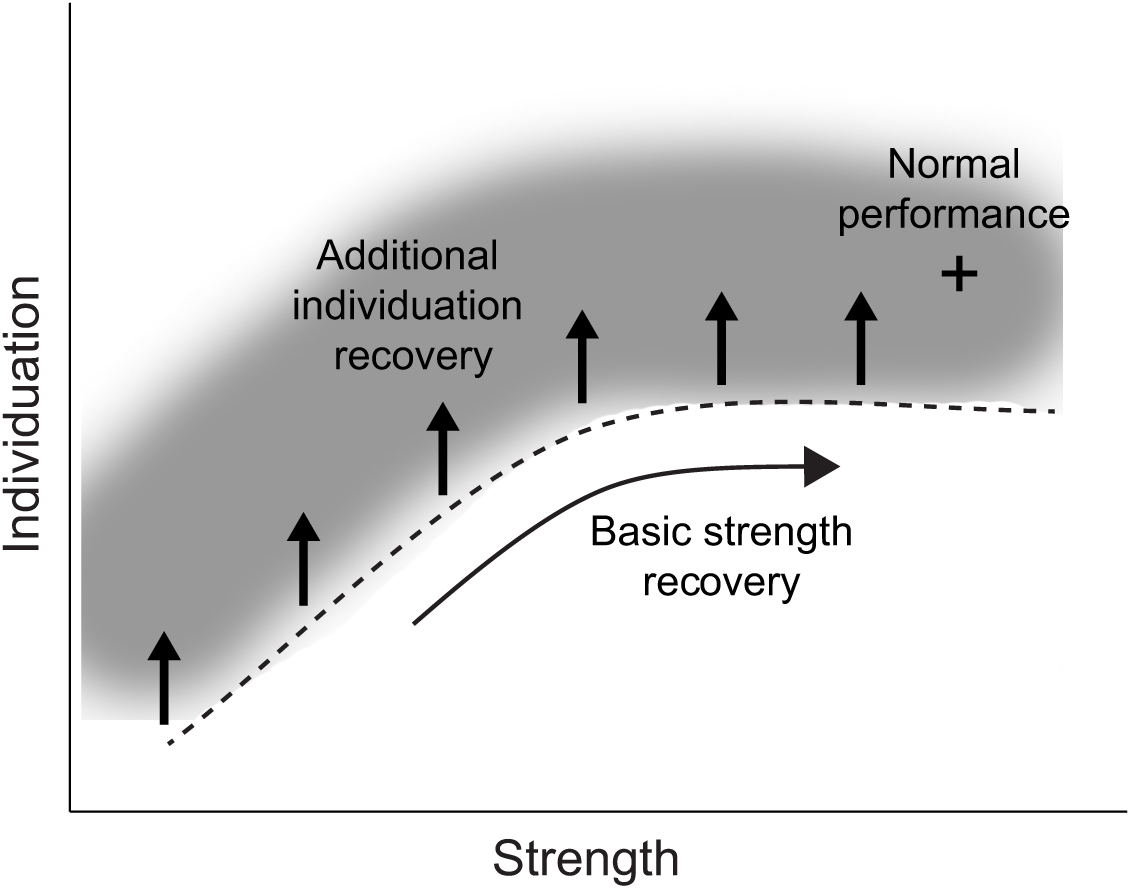
A schematic diagram of the hypothesis of two recovery systems. The first system (basic strength recovery) underlies strength recovery and a restricted amount of individuation recovery. This system therefore defines the lower bound (dashed line) of the space occupied by recovering patients (gray clound). A second system (additional inidividuation recovery) adds further individuation abilities on top of the basic strength recovery.

The second system would then add additional control capacities to the first system (vertical arrows in Fig. 6). Patients with a strong contribution from this second system may gain full recovery of individuation; patients with no or only partial contribution from the second system may recover completely in strength, but not in individuation. Importantly the recovery of this second system also occurs early after stroke, subsequently a patient’s relative position above or below the mean recovery function remains relatively fixed (Fig. 4D).

The lesion analysis adds support to the two-systems model for recovery suggested by the behavioral data. A wealth of evidence in humans and non-human primates implicates the role of CST in finger control, especially the monosynaptic corticomotoneuronal connections originating from “new” M1 (Lawrence and Kuypers, 1968; Porter and Lemon, 1993; Rathelot and Strick, 2009). Notably, these connections do not generate high levels of force but rather finely graded forces riding on top of larger forces (Maier *et al.*, 1993). Consistent with this idea, lesions in the gray matter of the hand areas in M1- the main origin of corticospinal projections- as well as the CST, correlated more with impaired individuation than with strength.

In contrast, finger strength may rely on other neural pathways, including the reticulospinal tract (RST), which can support strength and gross movements (Buford and Davidson, 2004; Davidson and Buford, 2004). Although the RST has been found to participate in some degree of finger control, its functional range is limited and biased towards flexor muscles (Riddle *et al.*, 2009; Baker, 2011).

Recovery after stroke is likely to result from the dynamic interplay between the CST and other descending pathways, particularly the RST. In this scenario, the correlation between strength and control at low levels of strength may represent the state of both the residual CST and of cortical projections to reticular nuclei in the brainstem. In this framework, recovery along the lower bound of the invariant function would represent the contribution of the RST and other non-CST descending pathways. Those patients with less damage to the CST would consistently ride above this function.

The dichotomy proposed here may be too simplistic. While the origin of the corticoreticular inputs is more diffuse (Keizer and Kuypers, 1989) and bi-laterally organized (Buford and Davidson, 2004; Sakai *et al.*, 2009; Soteropoulos *et al.*, 2012), many of the projections to the reticular formation arise from the primary motor cortex (Catsman-Berrevoets and Kuypers, 1976; Jones and Wise, 1977). Thus, our lesion ROIs will have included the corticoreticular tract to some degree, possibly explaining the lower, but nevertheless significant correlation with strength. Furthermore, it is very likely that direct corticospinal projections contribute to hand strength to some degree.

Interestingly, there was a small degree of impairment, especially in strength, in the hand ipsilesional to the stroke. This finding confirms previous reports of deficits in the non-paretic hand using clinical scales, e.g. muscle weakness measured by dynamometry (Colebatch and Gandevia, 1989), and dexterity measured with the Nine Hole Peg Test (9HPT) in (Noskin *et al.*, 2008). This ipsilesional impairment is consistent with positing a strength role for the RST because it projects bilaterally.

A limitation of the current study is that the paradigm is designed to assess weakness and enslaving in finger flexors, but not extensors. Because finger extensors play an important role in finger individuation, and have been particularly associated with the CST, it is possible that individuation in the extensors would also be more CST-dependent and the dual systems we are implying for the flexor might not apply to the same degree.

## Conclusions

Here we found that hand function after stroke can be partitioned into strength and strength-independent control. Most recovery of both these components occurred in the first three months after stroke, although strength continued to improve for up to six months. At any time point after stroke, strength and strength-independent control were related by an invariant curvilinear function: strength and some degree of control are correlated up to a certain strength level and then control saturates; some subjects showed additional improvement in individuation riding on top of the main recovery function. The results suggest that hand recovery is supported by two separable systems: one that mainly contributes to the generation of large forces, as in the power grip, and another that is responsible for more precise control of the digits at all levels of force. This behavioral and imaging evidence for two systems contributing to recovered hand function after stroke is consistent with the known characteristics of the CST and RST.

## Acknowledgement

We thank Adrian Haith and Martin Lindquist for helpful discussions about data analysis. This main study was supported by James S. McDonnell Foundation JMSF 220020220. Additional support came from a Scholar Award from the James S. McDonnell Foundation and a Grant from the Wellcome Trust (094874/Z/10/Z) to Jörn Diedrichsen. Andreas R. Luft is supported by the P&K Pühringer Foundation.

## References

Baker SN. The primate reticulospinal tract, hand function and functional recovery. J. Physiol. 2011; 589: 5603–5612.

Bates D, Mächler M, Bolker B, Walker S. Fitting Linear Mixed-Effects Models using lme4.

Brainard DH. The Psychophysics Toolbox. Spat. Vis. 1997; 10: 433–436.

Buford JA, Davidson AG. Movement-related and preparatory activity in the reticulospinal system of the monkey. Exp. Brain Res. 2004; 159: 284–300.

Catsman-Berrevoets CE, Kuypers HG. Cells of origin of cortical projections to dorsal column nuclei, spinal cord and bulbar medial reticular formation in the rhesus monkey. Neurosci. Lett. 1976; 3: 245–252.

Ceritoglu C, Oishi K, Li X, Chou M-C, Younes L, Albert M, et al. Multi-contrast large deformation diffeomorphic metric mapping for diffusion tensor imaging. NeuroImage 2009; 47: 618–627.

Colebatch JG, Gandevia SC. The Distribution of Muscular Weakness in Upper Motor Neuron Lesions Affecting the Arm. Brain 1989; 112: 749–763.

Dale AM, Fischl B, Sereno MI. Cortical surface-based analysis. I. Segmentation and surface reconstruction. NeuroImage 1999; 9: 179–194.

Davidson AG, Buford JA. Motor outputs from the primate reticular formation to shoulder muscles as revealed by stimulus-triggered averaging. J. Neurophysiol. 2004; 92: 83–95.

Duncan PW, Goldstein LB, Matchar D, Divine GW, Feussner J. Measurement of motor recovery after stroke. Outcome assessment and sample size requirements. Stroke J. Cereb. Circ. 1992; 23: 1084–1089.

Ellis MD, Sukal-Moulton T, Dewald JPA. Progressive shoulder abduction loading is a crucial element of arm rehabilitation in chronic stroke. Neurorehabil. Neural Repair 2009; 23: 862–869.

Fischl B, Rajendran N, Busa E, Augustinack J, Hinds O, Yeo BTT, et al. Cortical folding patterns and predicting cytoarchitecture. Cereb. Cortex N. Y. N 1991 2008; 18: 1973–1980.

Fisher RA. On the ‘Probable Error’ of a coefficient of correlation deduced from a small sample. Metron 1921: 1–32.

Fugl-Meyer AR, Jääskö L, Leyman I, Steglind S. The post-stroke hemiplegic patient. 1. a method for evaluation of physical performance. Scand. J. Rehabil. Med. 1975; 7: 13–31.

Guttman L. A basis for analyzing test-retest reliability. Psychometrika 1945; 10: 255–282.

Heller A, Wade DT, Wood VA, Sunderland A, Hewer RL, Ward E. Arm function after stroke: measurement and recovery over the first three months. J. Neurol. Neurosurg. Psychiatry 1987; 50: 714–719.

Holland PW, Welsch RE. Robust regression using iteratively reweighted least-squares. Commun. Stat. - Theory Methods 1977; 6: 813–827.

Jones EG, Wise SP. Size, laminar and columnar distribution of efferent cells in the sensory-motor cortex of monkeys. J. Comp. Neurol. 1977; 175: 391–438.

Jørgensen HS, Nakayama H, Raaschou HO, Vive-Larsen J, Støier M, Olsen TS. Outcome and time course of recovery in stroke. Part II: Time course of recovery. The copenhagen stroke study. Arch. Phys. Med. Rehabil. 1995; 76: 406–412.

Kamper DG, Harvey RL, Suresh S, Rymer WZ. Relative contributions of neural mechanisms versus muscle mechanics in promoting finger extension deficits following stroke. Muscle Nerve 2003; 28: 309–318.

Kamper DG, Rymer WZ. Impairment of voluntary control of finger motion following stroke: role of inappropriate muscle coactivation. Muscle Nerve 2001; 24: 673–681.

Keizer K, Kuypers HG. Distribution of corticospinal neurons with collaterals to the lower brain stem reticular formation in monkey (Macaca fascicularis). Exp. Brain Res. 1989; 74: 311–318.

Krakauer JW, Carmichael ST, Corbett D, Wittenberg GF. Getting neurorehabilitation right: what can be learned from animal models? Neurorehabil. Neural Repair 2012; 26: 923–931.

Laird NM, Ware JH. Random-effects models for longitudinal data. Biometrics 1982; 38: 963–974.

Lang CE, Schieber MH. Differential impairment of individuated finger movements in humans after damage to the motor cortex or the corticospinal tract. J. Neurophysiol. 2003; 90: 1160–1170.

Lang CE, Schieber MH. Reduced muscle selectivity during individuated finger movements in humans after damage to the motor cortex or corticospinal tract. J. Neurophysiol. 2004; 91: 1722–1733.

Li S, Latash ML, Yue GH, Siemionow V, Sahgal V. The effects of stroke and age on finger interaction in multi-finger force production tasks. Clin. Neurophysiol. Off. J. Int. Fed. Clin. Neurophysiol. 2003; 114: 1646–1655.

Li ZM, Latash ML, Zatsiorsky VM. Force sharing among fingers as a model of the redundancy problem. Exp. Brain Res. 1998; 119: 276–286.

Lyden PD, Lau GT. A critical appraisal of stroke evaluation and rating scales. Stroke J. Cereb. Circ. 1991; 22: 1345–1352.

Maier MA, Bennett KM, Hepp-Reymond MC, Lemon RN. Contribution of the monkey corticomotoneuronal system to the control of force in precision grip. J. Neurophysiol. 1993; 69: 772–785.

Mori S, Oishi K, Jiang H, Jiang L, Li X, Akhter K, et al. Stereotaxic white matter atlas based on diffusion tensor imaging in an ICBM template. NeuroImage 2008; 40: 570–582.

Noskin O, Krakauer JW, Lazar RM, Festa JR, Handy C, O’Brien KA, et al. Ipsilateral motor dysfunction from unilateral stroke: implications for the functional neuroanatomy of hemiparesis. J. Neurol. Neurosurg. Psychiatry 2008; 79: 401–406.

Oishi K, Faria A, Jiang H, Li X, Akhter K, Zhang J, et al. Atlas-based whole brain white matter analysis using large deformation diffeomorphic metric mapping: application to normal elderly and Alzheimer’s disease participants. NeuroImage 2009; 46: 486–499.

Oishi K, Faria AV, van Zijl PC, Mori S. MRI atlas of human white matter. Academic Press; 2010.

Oishi K, Zilles K, Amunts K, Faria A, Jiang H, Li X, et al. Human brain white matter atlas: identification and assignment of common anatomical structures in superficial white matter. NeuroImage 2008; 43: 447–457.

Oldfield RC. The assessment and analysis of handedness: the Edinburgh inventory. Neuropsychologia 1971; 9: 97–113.

Picard RR, Cook RD. Cross-Validation of Regression Models. J. Am. Stat. Assoc. 1984; 79: 575–583.

Raghavan P, Krakauer JW, Gordon AM. Impaired anticipatory control of fingertip forces in patients with a pure motor or sensorimotor lacunar syndrome. Brain 2006; 129: 1415–1425.

Rathelot J-A, Strick PL. Muscle representation in the macaque motor cortex: an anatomical perspective. Proc. Natl. Acad. Sci. U. S. A. 2006; 103: 8257–8262.

Rathelot J-A, Strick PL. Subdivisions of primary motor cortex based on cortico-motoneuronal cells. Proc. Natl. Acad. Sci. 2009; 106: 918–923.

Reinkensmeyer DJ, Lum PS, Lehman SL. Human control of a simple two-hand grasp. Biol. Cybern. 1992; 67: 553–564.

Riddle CN, Edgley SA, Baker SN. Direct and indirect connections with upper limb motoneurons from the primate reticulospinal tract. J. Neurosci. Off. J. Soc. Neurosci. 2009; 29: 4993–4999.

Sakai ST, Davidson AG, Buford JA. Reticulospinal neurons in the pontomedullary reticular formation of the monkey (Macaca fascicularis). Neuroscience 2009; 163: 1158–1170.

Schieber MH. Individuated finger movements of rhesus monkeys: a means of quantifying the independence of the digits. J. Neurophysiol. 1991; 65: 1381–1391.

Sharpless JW. The nine hole peg test of finger hand coordination for the hemiplegic patient. Mossmans Probl. Orientated Approach Stroke Rehabil. 1982

Soteropoulos DS, Williams ER, Baker SN. Cells in the monkey ponto-medullary reticular formation modulate their activity with slow finger movements. J. Physiol. 2012; 590: 4011–4027.

Sukal TM, Ellis MD, Dewald JPA. Shoulder abduction-induced reductions in reaching work area following hemiparetic stroke: neuroscientific implications. Exp. Brain Res. 2007; 183: 215–223.

Sunderland A, Tinson D, Bradley L, Hewer RL. Arm function after stroke. An evaluation of grip strength as a measure of recovery and a prognostic indicator. J. Neurol. Neurosurg. Psychiatry 1989; 52: 1267–1272.

Witham CL, Fisher KM, Edgley SA, Baker SN. Corticospinal Inputs to Primate Motoneurons Innervating the Forelimb from Two Divisions of Primary Motor Cortex and Area 3a. J. Neurosci. Off. J. Soc. Neurosci. 2016; 36: 2605–2616.

Yousry TA, Schmid UD, Alkadhi H, Schmidt D, Peraud A, Buettner A, et al. Localization of the motor hand area to a knob on the precentral gyrus. A new landmark. Brain J. Neurol. 1997; 120 (Pt 1): 141–157.

Zatsiorsky VM, Li ZM, Latash ML. Enslaving effects in multi-finger force production. Exp. Brain Res. 2000; 131: 187–195.

Zhang Y, Zhang J, Oishi K, Faria AV, Jiang H, Li X, et al. Atlas-guided tract reconstruction for automated and comprehensive examination of the white matter anatomy. NeuroImage 2010; 52: 1289–1301.

